# Dityrosine cross-link trapping of amyloid-β intermediates reveals that self-assembly is required for Aβ-induced cytotoxicity

**DOI:** 10.1101/2020.03.25.007690

**Authors:** Mahmoud B. Maina, Kurtis Mengham, Gunasekhar K. Burra, Youssra A. Al-Hilaly, Louise C. Serpell

## Abstract

Multiple chemical reactions, such as the production of reactive oxygen species (ROS) can lead to dityrosine (DiY) formation via the cross-linking of closely spaced tyrosine residues and this can serve as a marker for aging. Amyloid-β (Aβ) has been found to be DiY cross-linked in the brains of AD patients. *In vitro,* Aβ forms DiY cross-links via metal-catalysed oxidation (Cu^2+^ and H_2_0_2_) (MCO) leading to the formation of fibrils that are resistant to formic acid denaturation. However, copper is well known to influence and enhance self-assembly. Here, to investigate the interplay between self-assembly and DiY cross-linking we have utilised a non-assembly competent variant of Aβ (vAβ). MCO and UV oxidation experiments using vAβ and wild-type Aβ, revealed that DiY cross-linking stabilises, but does not induce or promote Aβ assembly. Cu^2+^ alone, without H_2_0_2_, facilitates the formation and DiY cross-linking of wild-type Aβ into long-lived oligomers. Our work reveals DiY formation halts further Aβ self-assembly. DiY cross-linked Aβ is non-toxic to neuroblastoma cells at all stages of self-assembly in contrast to oligomeric non-cross-linked Aβ. These findings point to a mechanism of toxicity that necessitates continuing self-assembly of the Aβ peptide, whereby trapped DiY Aβ assemblies and assembly incompetent variant Aβ are unable to result in cell death.

## Introduction

Alzheimer’s disease (AD) is the most common form of dementia, and it is characterised by the deposition of amyloid-β (Aβ) and Tau in extracellular plaques and intracellular neurofibrillary tangles, respectively. The amyloid cascade hypothesis implicates the pathological accumulation of Aβ and its aggregation from monomers into oligomers and fibrils, as a key event in the development of AD^1^. Many pieces of evidence have subsequently shown that the oligomeric form of Aβ is the most toxic species, resulting in a reformulated amyloid cascade hypothesis in which Aβ oligomers are proposed to be central to AD pathogenesis^2^. Indeed, accumulated evidence shows that Aβ oligomers disrupt cellular function in cultured cells and animal models^2–6^. Numerous studies have searched for the elusive “toxic” species and tried to characterise its structure. For example, I2mers, *56 KDa, hexamers have all be implicated as a specific structural species that interacts with particular receptors (e.g. NDMA etc) leading to downstream cell death.

Oxidative stress has been proposed as a key mechanism that mediates Aβ toxicity^7,8^. Indeed, it is one of the earliest sources of damage in human AD^9^. In a cellular model, we have also shown that oxidative stress is one of the earliest events induced by Aβ oligomers^10^. One of the ways through which oxidative stress causes cellular damage is through protein oxidation. The most common consequence of protein oxidation includes amino acid side chain modification, protein fragmentation and protein cross-linking (e.g. via dityrosine (DiY) crosslinking)^11^. DiY cross-linking is mediated via carbon-carbon bonding between two proximal tyrosines, resulting in stable, non-reversible covalent bond^12^. DiY cross-linking is known to provide elasticity, strength and stability to proteins and has been shown to form within proteins involved with neurodegenerative diseases (e.g. Aβ and a-synuclein)^13,14^. Indeed, we have previously shown the colocalisation of DiY with Aβ in plaques and α-synuclein in Lewy bodies in the human AD and Parkinson’s disease (PD) brain tissue, respectively^15,16^. This supports the potential disease relevance of DiY cross-linking in AD and PD.

To learn more about the importance of DiY cross-linking in AD, several studies have investigated the impact of DiY formation on Aβ aggregation and toxicity, mostly using metal-catalysed oxidation (e.g. Cu^2+^/H_2_0_2_) (MCO) and peroxidase-catalysed oxidation. Some studies have shown that DiY cross-linking of both Aβ40 and Aβ42 results in the generation of toxic Aβ assemblies with reduced assembly speed^17–22^, inhibit Aβ40 assembly, especially in highly oxidative environments^23^ or induce the formation of Aβ42 non-amyloidogenic aggregates when catalysed with a high concentration of Cu^2+^^24^. Whether DiY cross-linking is a driver, facilitator or inhibitor of Aβ self-assembly is still not clear from these studies. Moreover, given that Aβ self-assembly is rapid, the timepoint of the cross-linking during the assembly process may influence the nature of the cross-linked Aβ assemblies. Using MCO and UV-induced oxidation to induce DiY cross-linking, here we show that DiY formation results in the stabilisation of Aβ42 assemblies and prevents or very significantly slows further elongation of the assemblies. To specifically address the impact of DiY cross-linking on Aβ assembly, we compared the effect of oxidation on an non-assembly competent variant Aβ (vAβ)^3^ and revealed that DiY cross-linking does not induce or promote its assembly. We show that in the absence of H_2_0_2_, CuCl_2_ at a concentration found around Aβ plaques (~ 400 μM)^25^, is sufficient to facilitate the formation and cross-linking of Aβ42 oligomers into a long-lived oligomer population. Cell death assay revealed that unlike the self-assembling non-cross-linked Aβ, DiY-stabilised Aβ is non-toxic to neuron-like cells. Our results highlight the importance of self-assembly for Aβ toxicity and suggest that DiY cross-linking does not induce or promote Aβ assembly, instead, it stabilises Aβ assemblies.

## Materials and Methods

### Preparation of Aβ

Recombinant Aβ42 (Aβ) and variant Aβ42 (vAβ) were purchased in Hexafluoroisopropanol (HFIP) films from rPeptide and JPT, respectively. The peptides were prepared using an established protocol, and all procedures were done using protein LoBind Eppendorfs and tips^3^. 0.2 mg/mL of the peptides were solubilized in 200 μL HFIP (Sigma-Aldrich), vortexed for 1 min and sonicated in 50/60 Hz bath sonicator for 5 min. The HFIP was removed by air drying using a low stream of nitrogen gas. The dried peptide films were dissolved in 200 μL Dimethyl sulfoxide (DMSO) >99.9% (Sigma-Aldrich), vortex for 1 min and sonicated for 1 min. The solution was passed through a 2 mL 7K MWCO Zeba buffer-exchange column (Thermo Scientific) and stacked with 40 μL of 10 mM phosphate buffer (pH 7.4). The concentration of the peptides was determined using a NanoDrop spectrophotometer (Thermo Scientific) at a wavelength of 280 nm (extinction coefficient of 1490). The peptide solutions were immediately diluted to 50 μM with the 10 mM phosphate buffer and used as indicated. For the experiments involving preformed Aβ assemblies, the 50 μM Aβ was left to assemble for 24 without shaking before being subjected to UV exposure.

### Metal-catalysed oxidation (MCO) of Aβ and vAβ

Freshly prepared samples of Aβ and vAβ peptides (50 μM) in 10 mM phosphate buffer, pH7.4 were incubated i) without CuCl_2_, ii) in the presence of 400 μM CuCl_2_ (peptide: CuCl_2_ ratio 1:8) and iii) in the presence of 400 μM CuCl_2_ and 2.5 mM H_2_0_2_. The peptides were incubated at 37°C without agitation and at each time point collected, the oxidation reaction was stopped using ethylenediaminetetraacetic acid (EDTA) at a final concentration of 2 mM. A minimum of three independent experiments were conducted to ensure the reproducibility of the findings.

### Photo-oxidation of Aβ and vAβ

Freshly prepared samples of Aβ and vAβ peptides (50 μM) in 10 mM phosphate buffer, pH 7.4 were incubated i) without UV-C in the dark, and ii) under of UV-C for 5min or 2h using a G6T5 Germicidal 9’ 6W T5 UVC lamp set to 8 J/m^2^/sec (General Lamps Ltd). A minimum of three independent experiments were conducted to ensure the reproducibility of the findings.

### Fluorescence spectroscopy

The formation of dityrosine was monitored with a fluorescence spectrophotometer (Varian Ltd., Oxford, UK), using a 1 cm path length quartz cuvette (Starna, Essex, UK). The presence of dityrosine was detected using fluorescent excitation wavelength of 320 nm and emission collected between 340 – 600 nm, with dityrosine peak signal observed between 400-420 nm. Tyrosine fluorescence signal was monitored using an excitation wavelength of 280 nm and an emission wavelength of 305 nm, with the peak tyrosine emission observed at 305 nm. For experiments involving metal-catalysed oxidation, the reaction was stopped using EDTA to a final concentration of 2 mM. For all the measurements, the excitation and emission slits were both set to 10 nm, scan rate set to 300 nm/min with 2.5 nm data intervals and an averaging time of 0.5 s. The photomultiplier tube detector voltage was set at 500 V.

### Thioflavin T fluorescence assay of Aβ self-assembly

Samples were incubated with 100 μM Thioflavin T (Th-T), and the rate of Th-T binding was monitored over time at 37°C using SpectraMax i3 plate reader. The readings were collected in a black 96-well plate with a clear bottom (PerkinElmer), which was sealed with an optically clear polyolefin film to avoid evaporation (StarSeal Advanced Polyolefin Film, Starlab). The number of reading per well was set to 6, PMT voltage was set to high and blank spectra of the buffer were subtracted to protein fluorescence scans. The excitation wavelength was set at 440 nm, and emission at 483 nm and the signal collected every 30 min, with 5 sec low orbital shakes before readings. The fluorescence data were plotted against time. A minimum of three independent experiments was repeated to ensure the reproducibility of the findings.

### Circular Dichroism (CD)

The secondary structure of Aβ and vAβ peptides at 50 μM concentration in 10 mM phosphate buffer (pH 7.4) incubated under different conditions was assessed using Jasco J715 CD spectrometer (Jasco, Goh-Umstadt, Germany). 140 μL of each sample was placed into a 1 mm path length quartz cuvette (Hellma) and scanned between 190 nm and 260 nm. The CD spectra were collected in triplicate at a maintained temperature of 21 °C.

### Negative-stain transmission electron microscopy (TEM)

The morphology of control and cross-linked Aβ and vAβ peptides was assessed by negative stain TEM. Briefly, 4 μL of each sample was dropped onto 400-mesh carbon-coated grids (Agar Scientific, Essex, UK). After 1 min incubation, the excess sample was blotted using filter paper, and the grid was washed with 4 μL filtered Milli-Q water and blotted. The grid was then negatively stained for 40 sec using 4 μL of filtered 2% (w/v) uranyl acetate. The excess stain was blotted with filter paper and grids left to air-dry before storage. The grids were examined on a Jeol Jem1400-plus transmission electron microscope (Jeol, USA), operated at 80 kV fitted with a Gatan Orius SC100 camera (UK).

### Dot blotting

A total of 4 μl was spotted onto a 0.2 μM pore nitrocellulose membrane and allowed to dry for 10 min. The membrane was boiled with PBS for 1 min twice and then blocked with blocking buffer (5% milk in 0.05% TBS-T) for 1 hour at room temperature on a rocker. The blocking buffer was next replaced with mouse NU-1 primary antibody (1/2000) and left to bind overnight at 4°C on a rocker. The membrane was washed 6 times for 5 min with washing buffer (0.05% TBS-T), then incubated with an HRP-conjugated goat anti-mouse secondary antibody for 1 hour. The membrane was washed six times for 5 min with washing buffer, then incubated with Clarity Western ECL Substrate (Bio-Rad) for 1 min before being developed in the darkroom. The NU-1 antibody was a gift from the William Klein lab^26^. A minimum of three independent experiments were conducted to ensure the reproducibility of the findings.

### Cell death assay

Differentiated SHSY5Y neuroblastoma cells were used for the toxicity experiments. Firstly, undifferentiated SHSY5Y neuroblastoma cells (ATCC CRL-2266™), were maintained in Dulbecco’s Modified Eagle Medium: Nutrient Mixture F-12 (DMEM/F-12) (Life Technologies, United Kingdom), supplemented with 1% (v/v) L-glutamate (L-Gln) (Invitrogen), 1% (v/v) penicillin/streptomycin (Pen/Strep) (Invitrogen) and 10% (v/v) Fetal Calf Serum at 37°C and 5% CO_2_. The undifferentiated SHSY5Y cells were seeded to 60% confluency in a CellCarrier-96 Ultra Microplates (PerkinElmer). The cells were differentiated in a medium containing 1% Fetal Calf Serum supplemented with 10 μM trans-Retinoic acid (Abcam) for 5 days. Next, the medium was replaced with a serum-free media supplemented with 2 nM brain-derived neurotrophic factor (BDNF) (Merck Millipore). After 2 days in the BDNF-containing media, the media was replaced with serum-free media and the cells were treated with UV cross-linked or uncross-linked vAβ or Aβ for 3days. At the end of the incubation period, the cells were incubated with ReadyProbes reagent (Life Technologies) for 15 min. The ReadyProbes kit contains NucBlue Live reagent that stains the nuclei of all live cells and Propidium iodide that stains the nuclei of dead cells with compromised plasma membrane. The cells were imaged at 37°C and 5% CO_2_ using Operetta CLS high-content analysis system (PerkinElmer) using DAPI and TRITC filters. At least 5000 dead and live cells were analysed using the Harmony software automated analysis algorithm within the Operetta CLS high-content analysis system. A minimum of three independent experiments were repeated to ensure the reproducibility of the findings.

## Results

### *In vitro* metal-catalysed oxidation results in the formation of dityrosine on wild-type and vAβ peptides

To investigate the influence of DiY cross-linking on Aβ assembly, we compared the effect of metal-catalysed oxidation (MCO) using CuCl_2_ and H_2_0_2_, on wild-type Aβ42 and variant Aβ42 (henceforth called Aβ and vAβ, respectively). vAβ is a variant of Aβ1-42 with F19S and G37D mutations which render the peptide assembly incompetent (see^3^ for detail).

We have previously detected DiY cross-links in Aβ and a-synuclein using a combination of techniques and shown that DiY can reliably be detected using fluorescence spectroscopy^15,16^. Here, rapid formation of DiY was detected for Aβ and vAβ samples incubated with both CuCl_2_/H_2_0_2_ (Aβ/CuCl_2_/H_2_0_2_ and vAβ/CuCl_2_/H_2_0_2_ – henceforth called MCO reaction), showing a fluorescence peak at 410 nm after only 5 mins, while CuCl_2_ alone or buffer only samples showed no peak (Fig. 1A). Detection of Tyrosine with an excitation/emission of 280/305 nm showed that the formation of DiY was matched by a decrease in tyrosine fluorescence in both the Aβ and vAβ MCO reactions compared to the samples incubated with CuCl_2_ alone or buffer only (Fig. 1B). DiY continued to increase for the MCO reactions upto 2h but did not increase further after 5 days (Fig 1C). However, by 5 days, small peaks could be observed for samples incubated with CuCl_2_ alone (Fig. 1D) although the DiY signal remained negligible for buffer only samples after 5 days incubation in the dark. Aβ and vAβ with CuCl_2_ samples showed a decrease in tyrosine fluorescence as early as 5 min of incubation with CuCl_2_ (Fig. 1B), indicating that the CuCl_2_ rapidly induces conformational changes in both Aβ and vAβ, independent of DiY cross-linking, which only starts appearing ~2h post-incubation. Overall, this suggests that the incubation of both Aβ and vAβ with CuCl_2_ alone or in combination with H_2_0_2_ induces DiY cross-linking, though at different speeds, suggesting that the timing of the DiY formation may differentially impact Aβ assembly properties.

**Figure 1.**
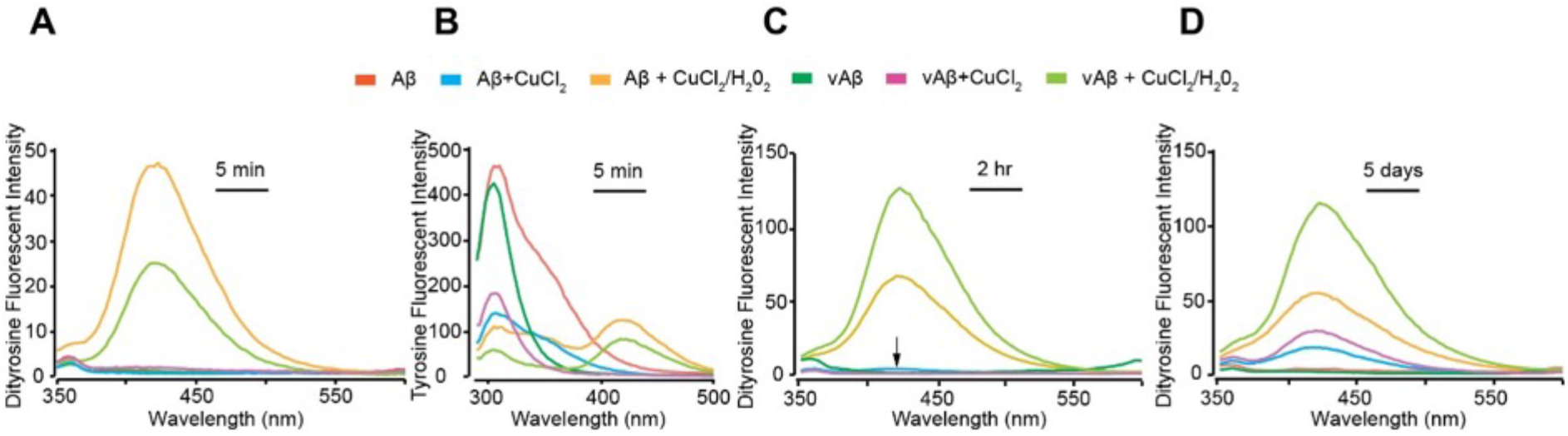
DiY formation in Aβ and vAβ via metal-catalysed oxidation. Freshly prepared Aβ and vAβ (50 μM) were incubated at 37°C, without CuCl_2_, with CuCl_2_ (400 μM) or CuCl_2_ in combination with H_2_0_2_. Fluorescence spectra were collected 5 min post-incubation using fluorescent excitation wavelength of 320 and emission collected between 340 – 600 nm, with DiY peak signal observed between 400-420 nm **(A)**. Fluoresence spectra were also collected at 5 min using an excitation wavelength of 280 nm and emission collected between 290 – 500, with peak tyrosine signal observed at 305 nm **(B)**. Fluoresence spectra were collected again at 2h **(C)**, and then 5 days **(D)** to follow DiY formation over time. A minimum of three independent experiments was repeated to ensure the reproducibility of the findings.

### The timing of dityrosine cross-linking influences the assembly of wild-type Aβ, but does not impact on the structure and aggregation propensity of variant Aβ

We have previously shown that DiY forms in both Aβ oligomers and fibrils^15^. Thioflavin T (Th-T) fluorescence assay was used to investigate whether DiY formation influences the Aβ assembly rate. As expected, the assembly incompetent vAβ incubated in buffer shows no increase in Th-T fluorescence with time^3^. Wild-type Aβ gave the expected Th-T spectra showing a sigmoidal curve (Fig. 2A). However, the Aβ and vAβ MCO reactions showed no Th-T fluorescence signal increase over the time frame of 50 h, indicating either no self-assembly or that the assembly is significantly slow that the threshold of Th-T detection has not been reached at the time points studied. Wild-type Aβ incubated with CuCl_2_ shows a small fluorescence signal for DiY at ~ 2h (Fig. 1C) and ThT spectrum showed a short lag phase (approximately up to 2h) and plateaus at a low fluorescence signal. This appears to suggest that the Aβ assemblies formed become stabilised without further elongation (Fig. 2A). vAβ CuCl_2_ showed no increase in Th-T fluorescence consistent with the absence of assembly under these conditions.

**Figure 2.**
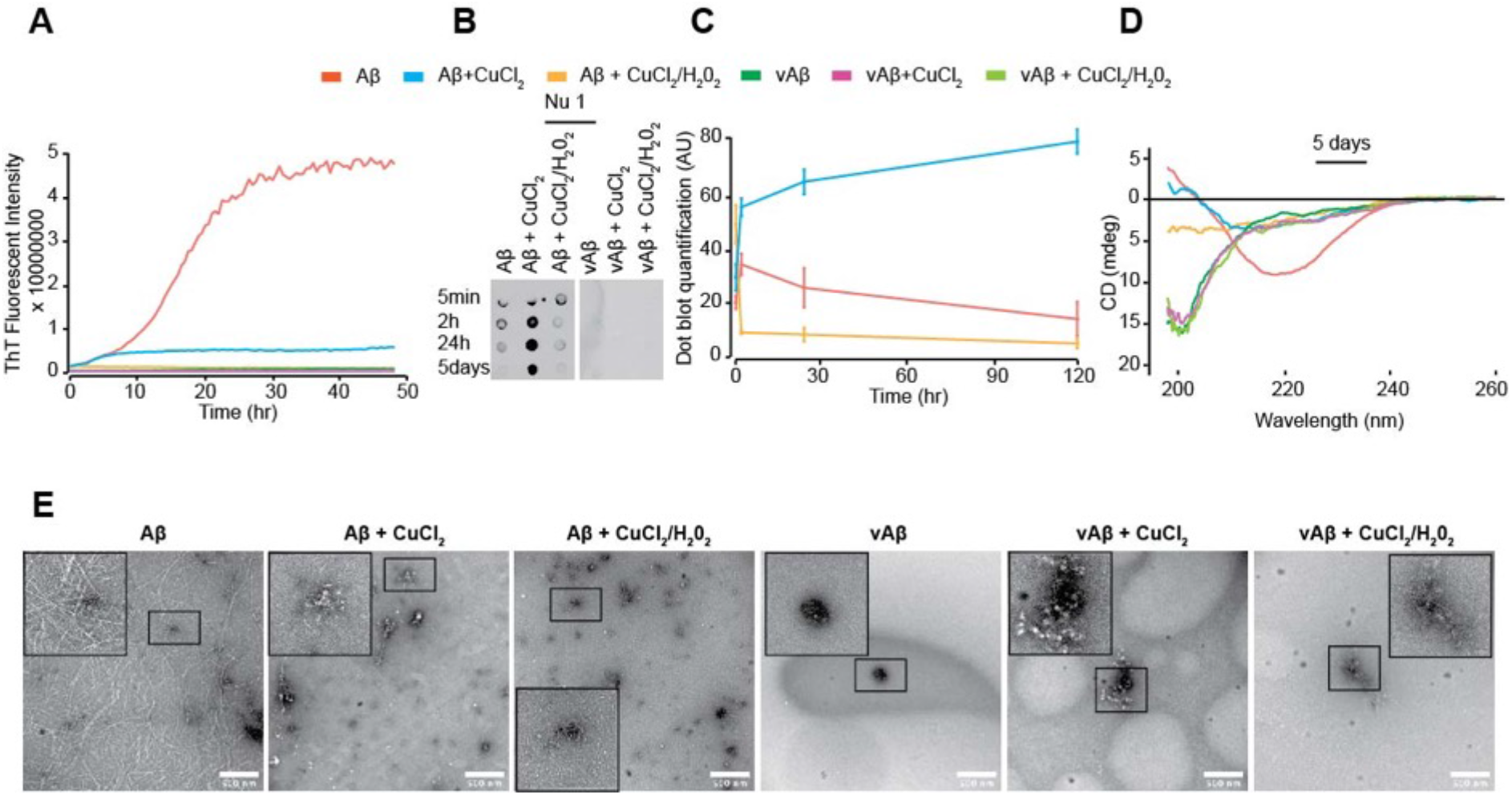
DiY cross-linking by metal-catalysed oxidation stabilises Aβ assemblies. Th-T fluorescence was monitored for the freshly prepared Aβ and vAβ (50 μM) incubated at 37°C, without CuCl_2_, with CuCl_2_ (400 μM) or CuCl_2_ in combination with H_2_0_2_ (**A**). Dot blotting using NU-1 antibody identified Aβ oligomers in the Aβ samples but not in vAβ reactions (**B**). Quantification of dot blotting signal against time reveals that the oligomers in the CuCl_2_-cross-linked Aβ remain stable for 5days, unlike in the other reactions which very low NU1 affinity signal (**C**). CD at 5 days showed a high β-sheet content in the uncross-linked Aβ reaction, with reduced signal for CuCl_2_ Aβ and CuCl_2_/H_2_0_2_ Aβ. All vAβ samples showed spectra indicating random coil conformation (**D**). TEM imaging at 5 days revealed a network of fibres in the uncross-linked Aβ, while the CuCl_2_ Aβ showed clumped assemblies with very little fibre density and the CuCl_2_/H_2_0_2_ revealed amorphous-like assemblies. Uncross-linked vAβ showed no assemblies while both cross-linked vAβ reactions showed amorphous-like aggregates (**E**). A minimum of three independent experiments was repeated to ensure the reproducibility of the findings.

We have previously shown that our Aβ preparation method results in the generation of monomers that aggregate into oligomers detected by the oligomer specific antibody, NU-1^26^, before forming fibrils and amyloid plaques^3^. As expected, dot blotting with the NU-1 antibody revealed the presence of oligomers at 2h in buffer incubated Aβ samples, which disappeared over time (Fig. 2B). In contrast, the buffer incubated vAβ reactions showed no NU-1 reactivity, indicating the absence of oligomer formation as previously reported (Fig. 1F)^3^. Aβ oligomers were only minimally detected in the Aβ MCO reaction at 5 min, which disappeared over time (Fig. 2B, C). No NU-1 reactivity was observed for vAβ MCO reaction at any of the time points measured. Interestingly, the Aβ CuCl_2_-cross-linked reaction formed more oligomers early-on, which persisted throughout the time studied, suggesting that slower cross-linking by CuCl_2_ facilitates Aβ oligomer formation and stabilisation via DiY cross-linking and traps Aβ in a NU-1 affinity conformation (Fig. 2B, C).

Circular Dichroism (CD) and Transmission Electron Microscopy (TEM) were used to study the secondary structure and morphologies of the resulting assemblies after five days incubation. Spectra from vAβ under all three conditions showed a trough at 198nm consistent with random coil conformation. Aβ in buffer showed a minimum at 218 nm consistent with an expected β-sheet conformation, typical of fibrillar Aβ (Fig. 2D). Aβ CuCl_2_ shows a broad but weak minimum at 218 nm, but the signal from Aβ MCO reaction is too weak for assignment of secondary structure. This may be consistent with loss of protein from solution.

Electron micrographs showed an extensive fibril network for buffer incubated Aβ while CuCl_2_-cross-linked Aβ showed clumped assemblies with a significantly reduced fibre density, and the MCO Aβ reaction revealed amorphous structures with very few or no fibres (Fig. 2E). Buffer incubated vAβ sample showed very few assemblies of any kind. The vAβ MCO and CuCl_2_ reactions exhibited a few amorphous-like clumped assemblies. The CD and TEM data suggest that the MCO catalysed DiY cross-linking of Aβ prevents the formation of β-sheet-rich amyloid fibrils, while CuCl_2_ catalysed DiY cross-linking generates oligomeric Aβ assemblies with some β-sheet conformation. Taken together, the results from Th-T, dot blotting, CD and TEM revealed that the rapid DiY cross-linking of Aβ induced through MCO results in the association and trapping of Aβ as intermediates at the stage of assembly and inhibit their further assembly, while CuCl_2_ facilitates the formation and DiY cross-linking of Aβ oligomers into long-lived oligomer population. In contrast, none of the conditions induced any assembly for vAβ into amyloid fibrils. Instead, the MCO and CuCl_2_ reactions resulted in association and trapping of of vAβ into amorphous assemblies that show no evidence of β-sheet conformation.

### Photo-oxidation induces dityrosine crosslinking on wild-type Aβ and vAβ peptides

The results above suggest that copper contributes to DiY formation but previous studies have suggested that metals influence the assembly of Aβ^18,23,24,27^. Therefore, to avoid the complication of using metals which may influence assembly, UV photo-oxidation was used to oxidise Aβ and vAβ (Fig. 3). Fluorescence spectra showed a small peak at 410 for both Aβ and vAβ after only 5 mins incubation in UV. However, the intensity was lower than under MCO conditions (Fig. 3A). After 2h of UV exposure, a significant increase in intensity at 410 nm was observed (Fig. 3B). Following 2h of UV exposure, samples were stored in the dark. However, the intensity continued to increase for 120h following incubation (Fig. 3C). These results show that UV photo-oxidation can efficiently induce DiY cross-linking for Aβ and vAβ peptides. The results are compared to peptides incubated without UV exposure for reference. We can not rule out the oxidation of other residues by the UV irradiation (e.g Met35), but here we focus on DiY.

**Figure 3.**
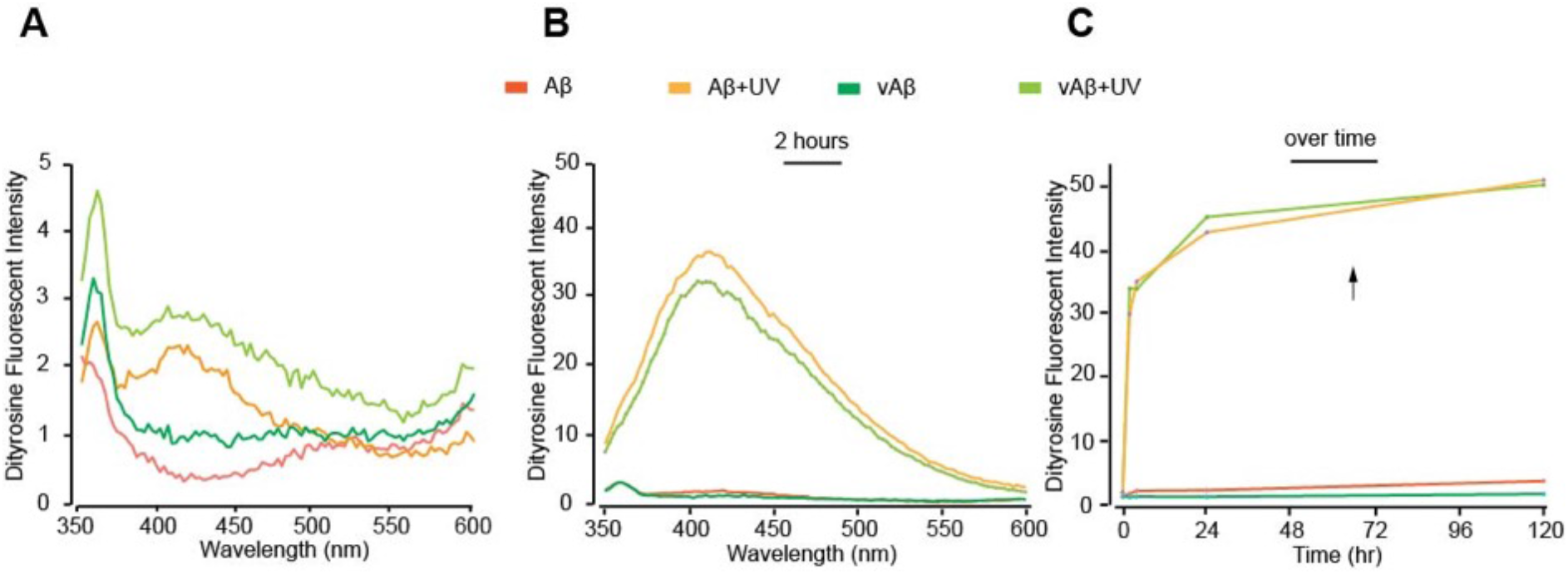
DiY formation in Aβ and vAβ via UV photo-oxidation. Freshly prepared Aβ and vAβ (50 μM) was incubated under UV. Fluorescence spectra were collected 5 min post-incubation using fluorescent excitation wavelength of 320 and emission collected between 340 – 600 nm, with DiY peak signal observed between 400-420 nm after 5 min of incubation (**A**), which increased following 2h of incubation (**B**). Fluorescence intensity at 410 nm against time showed that incubation of the 2h UV-exposed Aβ and vAβ samples in the dark resulted in futher increase in DiY formation in the absence of the UV (**C**). The Aβ and vAβ samples that were not exposed to UV showed no DiY signal. A minimum of three independent experiments was repeated to ensure the reproducibility of the findings.

### DIY crosslinking by photo-oxidisation influences the assembly of wild-type Aβ, but does not impact on the structure and assembly of vAβ

Th-T fluorescence was used to monitor assembly of Aβ and vAβ following UV photo-oxidation for 2 hours. The results are compared to peptides incubated without UV exposure for reference. A small increase in Th-T fluorescence was observed for Aβ+UV after about 25h incubation, while vAb+UV showed no fluorescence intensity (Fig. 4A). NU-1 dotblots again showed no reactivity to vAb+UV (Fig. 4B). The Aβ+UV sample showed lower level NU-1 reactivity at 2h compared to the Aβ control (Fig. 4B), and reactivity disappeared after 5 days post-UV incubation in the cross-linked Aβ+UV sample, even though a small quantity of oligomers could still be detected in the control Aβ sample (Fig. 4B). CD showed spectra consistent with random coil conformation for both vAβ and Aβ incubated under UV and a loss of signal which was more evident for Aβ than for vAβ (Fig. 4C). By TEM, Aβ+UV and vAβ+UV again showed small assemblies at 2h, which are still present alongside, occasionally clumped amorphous-like assemblies after five days. Similar to the results from MCO DiY cross-linking (Fig. 2B), this suggest that the UV-induced DiY cross-linking leads to the stabilisation of Aβ and vAβ in trapped and in an assembly incompetent species.

**Figure 4.**
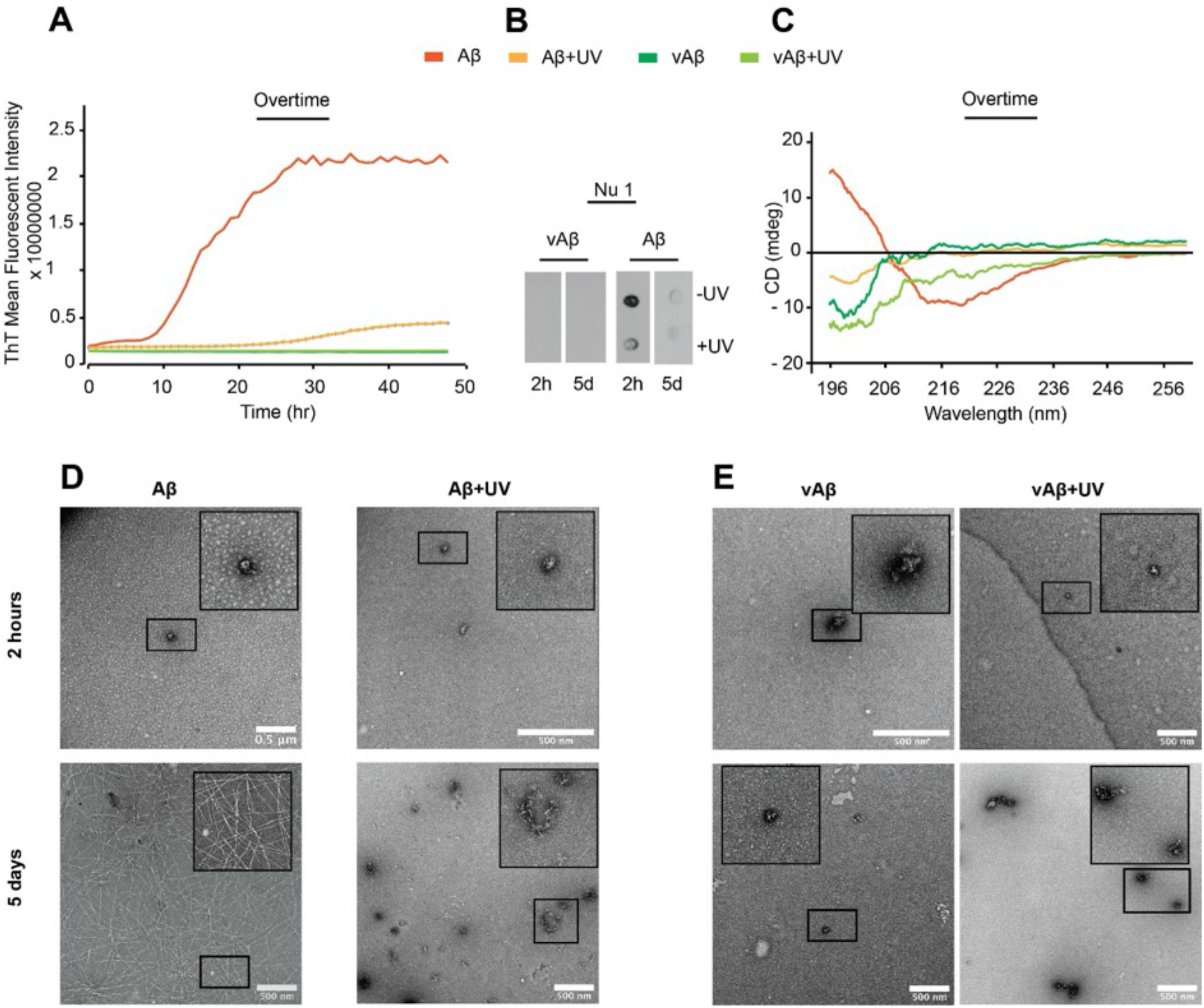
UV-induced DiY cross-linking of early Aβ assemblies prevents further elongation. (**A**) Th-T fluorescence spectrum show the expected increase in fluorescence for assembling Aβ but Aβ+UV ThT fluorescence did not increase with time. vAβ incubated in the absence or presence UV showed no Th-T binding. (**B**) Dot blotting using NU-1 antibody shows binding suggesting fewer oligomers in the cross-linked Aβ+UV than in the uncross-linked Aβ sample. No binding of NU1 was observed for Aβ+UV 5 days post-UV exposure, but a small signal was detected in the Aβ-UV sample. (**C**) CD at 5 days showed a high β-sheet content in the Aβ-UV sample, while the cross-linked Aβ+UV showed a loss of signal but indicting some random coil. Cross-linked and uncross-linked vAβ samples showed random coil conformation. (**D-E**) TEM after 2h and 5 days showed that the uncross-linked Aβ at 2h formed oligomers, which transformed into a network of fibres at 5 days. The cross-linked Aβ+UV samples formed small assemblies at 2h, some of which developed into amorphous-like assemblies at 5 days. vAβ does not assemble into amyloid fibrils but vAβ+UV forms some amorphous aggregates after 5 days. A minimum of three independent experiments was repeated to ensure the reproducibility of the findings.

### Co-incubation with DIY crosslinked Aβ slows the assembly of freshly prepared Aβ

Our findings thus far suggest that DiY cross-linking stabilise Aβ assemblies and prevent them from further elongation into amyloid fibrils. To investigate this further, DiY cross-linking was induced in Aβ using 2h UV exposure, and the sample was then incubated with an equal concentration of freshly prepared Aβ (20 μM:20 μM) and compared to a 40 μM Aβ sample not exposed to UV. Fluorescence spectroscopy confirms the presence of DiY in the 40 μM Aβ sample exposed to UV and the signal reduced by half when the cross-linked 40 μM Aβ was diluted to 20 μM (Fig. 5A). 20 μM cross-linked Aβ added to 20 μM freshly prepared uncrosslinked Aβ (Aβ+UV:Aβ), revealed the presence of DiY signal (Fig. 5A). Interestingly, the levels of DiY in the Aβ+UV:Aβ mixture was higher than that of 20 μM cross-linked Aβ, suggesting that the presence of the UV incubated Aβ in the environment results in the cross-linking of the freshly prepared Aβ following the co-incubation. Th-T fluorescence showed that the 40 μM Aβ sample showed a shorter lag phase and higher Th-T signal compared to the 20 μM Aβ sample. The 20 μM and 40 μM cross-linked Aβ samples showed no Th-T binding, similar to previous observations. When the Aβ mixtures were incubated together (Aβ+UV:Aβ 1:1), the mixture showed a longer lag phase (+20h) and lower intensity Th-T signal than the 20 μM uncrosslinked Aβ even after 50h (Fig. 5B). This suggests inhibition of self-assembly in the Aβ+UV:Aβ-UV mixture. At 4 days, TEM imaging revealed the presence of Aβ fibrils in both 20 μM and 40 μM Aβ samples, which were not detected in the cross-linked samples (Fig. 5C). Only scant fibrils and smaller Aβ assemblies could be detected in the Aβ+UV:Aβ mixture, further confirming a reduced assembly in this mixture (Fig. 5C). Taken together, this suggests that the cross-linking of freshly prepared Aβ induced by the DiY cross-linked Aβ environment, leads to the stabilisation of some Aβ assemblies, resulting in the slower aggregation kinetics observed and prevention of elongation. This also suggests that the DiY species may bind to Aβ and inhibit further elongation.

**Figure 5.**
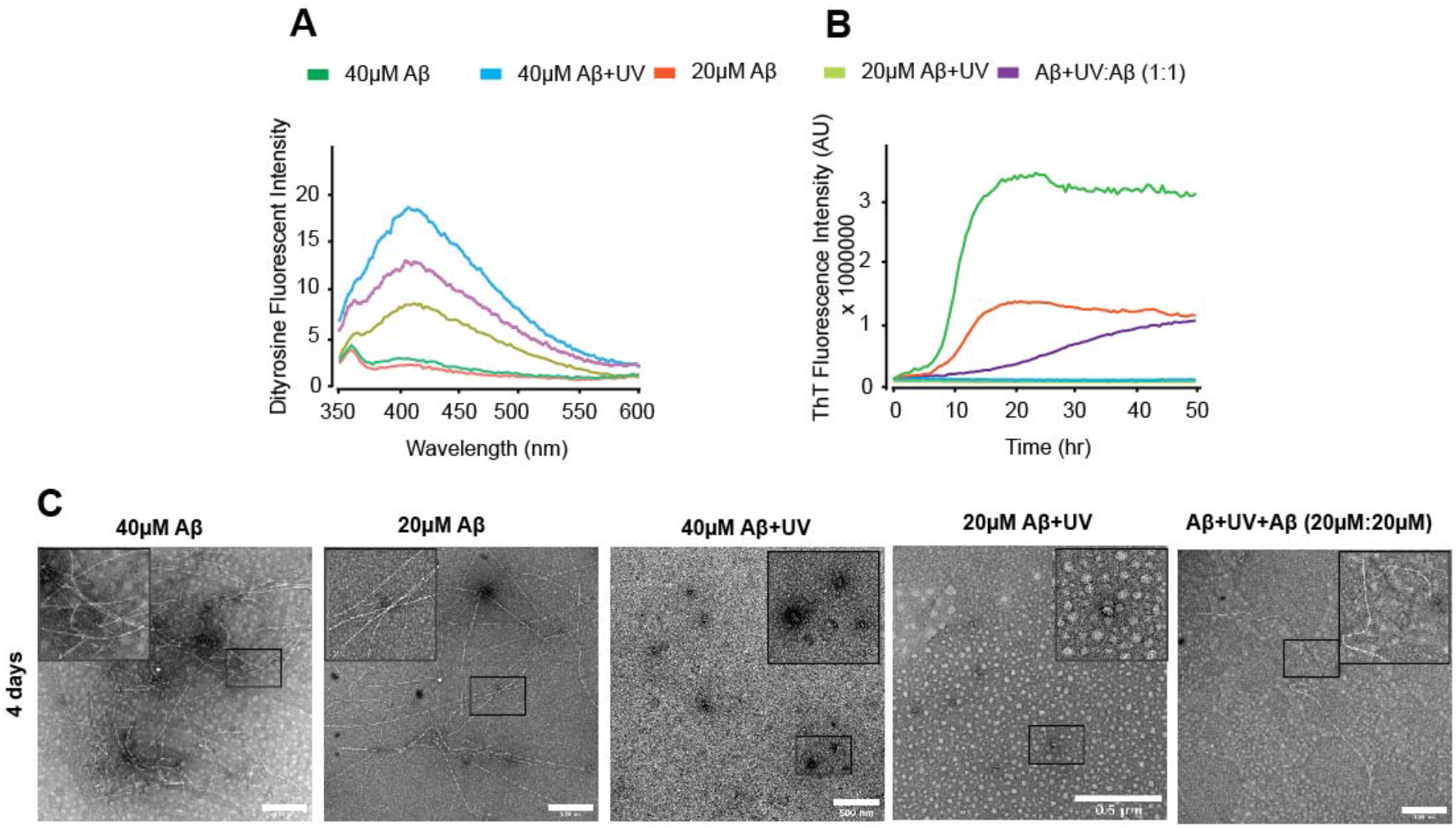
Co-incubation with DiY cross-linked Aβ assemblies slows the aggregation of freshly prepared Aβ. (**A**) DiY fluorescence was measured for Aβ (40 and 20 μM) and Aβ+UV:Aβ (20 μM: 20 μM) mixture formed from co-incubation of freshly prepared Aβ with DiY-cross-linked Aβ. DiY signal was not detected in uncross-linked freshly prepared 40 and 20 μM Aβ but was induced in Aβ+UV samples and in the mixture. (**B**) Th-T fluorescence showed that the uncross-linked 40 and 20 μM Aβ assemble at different rates, which was significantly delayed for Aβ+UV:Aβ-UV mixture and completely absent in the cross-linked 40 and 20 μM Aβ samples. (**C**) TEM imaging at 4 days revealed that the presence of fibrils in both uncross-linked 40 and 20 μM Aβ, which was significantly reduced in the Aβ+UV:Aβ-UV mixture, and absent in the cross-linked 40 and 20 μM Aβ samples. A minimum of three independent experiments was repeated to ensure the reproducibility of the findings.

### Cross-linking of pre-formed Aβ assemblies slows further assembly of Aβ oligomers/protofibrils and prolongs their half-life

Our findings thus far strongly suggest that DiY crosslinking stabilise and strongly slows the aggregation and further elongation of Aβ assemblies. However, it is not clear whether DiY cross-linking could also alter the assembly of preformed Aβ assemblies. To investigate this, Aβ (50 μM) was freshly prepared and allowed to assemble for 24h (henceforth called aAβ). aAβ samples were exposed to 2h UV to induce DiY cross-linking (aAβ+UV) (Fig. 6). Unlike samples that were not exposed to UV (aAβ), the aAβ+UV samples showed an increasing intensity arising from DiY, which continued to form even after incubation in the dark following the UV exposure (Fig. 6A). Th-T fluorescence intensity showed that aAβ continues to assemble, reaching plateau after about 15h (Fig. 6B). However, DiY cross-linking induced by UV leads to the inhibition of further assembly of the aAβ+UV for ~ 40h, suggesting that the cross-linking leads to the stabilisation or trapping of aAβ+UV (Fig. 6B). A gradual increase in fluorescence is observed beyond 40h which may indicate delayed assembly. Dot blotting using NU1 revealed a comparable level of oligomers in the cross-linked and uncross-linked aAβ samples (Fig. 6C). Interestingly, the oligomers in the cross-linked aAβ+UV sample persisted beyond 4days, unlike the uncross-linked aAβ-UV sample which showed a very low level of oligomers at the later time point (Fig. 6C). TEM imaging showed the presence of oligomeric assemblies and scant fibres in the cross-linked aAβ+UV sample, compared to the uncrosslinked aAβ-UV which showed extensive fibril network. Together, this data provides further evidence indicating that DiY crosslinking leads to the stabilisation of aAβ assemblies to prevent or delay further elongation. This is similar to the stabilisation of Aβ oligomers observed following the slower/milder DiY cross-linking in the Aβ/CuCl_2_ preparation which occurred after the formation of Aβ oligomers (Fig. 2B, C).

**Figure 6.**
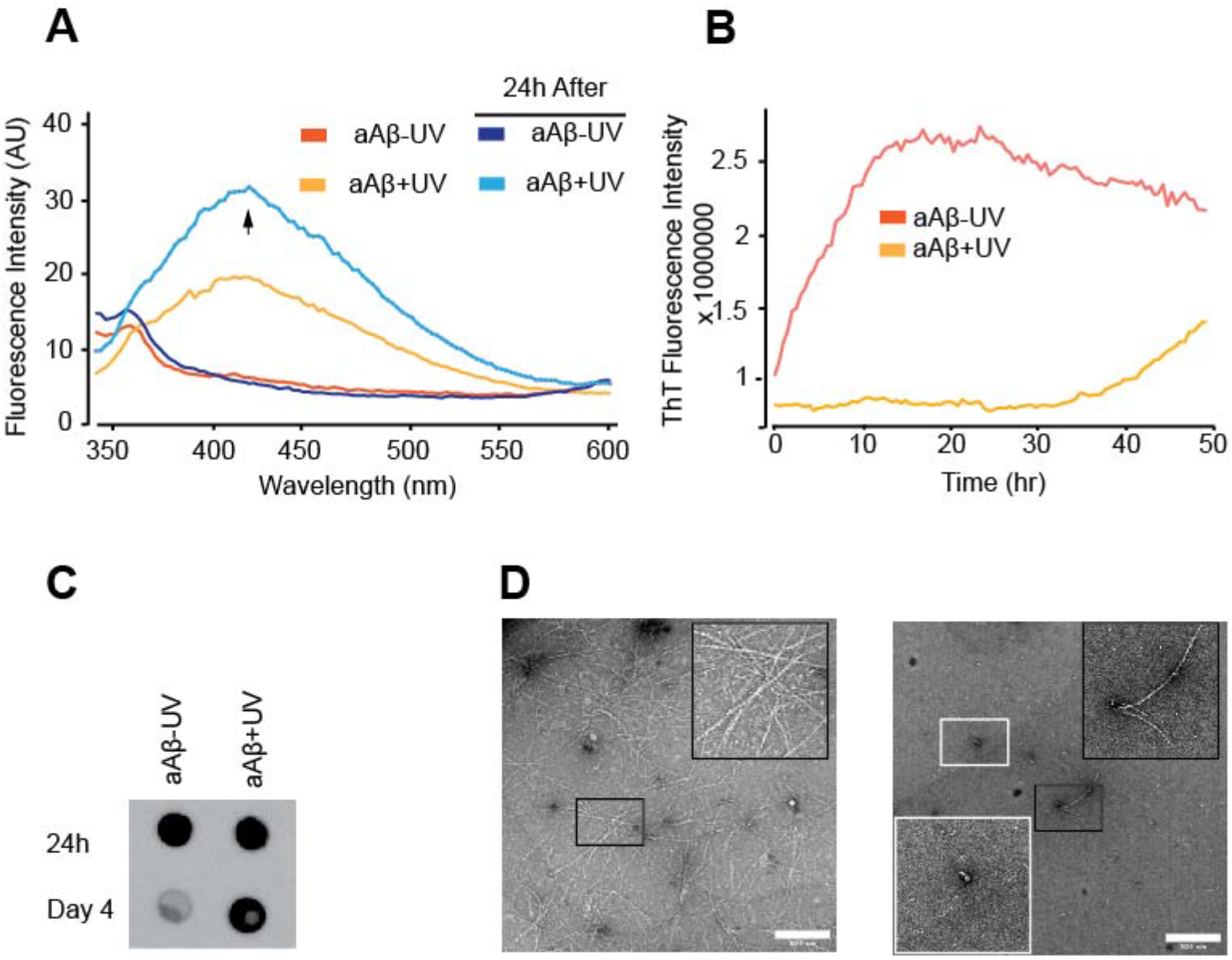
Cross-linking of pre-formed Aβ (aAβ) (24h) assemblies slows further assembly of Aβ oligomers/protofibrils and prolongs the half-life of the oligomers. (**A**) DiY signal was detected in the cross-linked aAβ+UV sample, which continued over time. Uncross-linked aAβ-UV samples showed no DiY signal. (**B**) Th-T fluorescence showed that the uncross-linked aAβ-UV assemblies continue to assemble, while the cross-linked aAβ+UV became significantly inhibited from further elongation upto 40 h. (**B**) Dot blotting using NU-1 antibody reveals the presence of oligomers in the uncross-linked and uncross-linked aAβ at the starting time point. However, the oligomers are still strongly detected in the cross-linked aAβ+UV at 4 days, unlike in the uncross-linked samples. (**D**) TEM imaging at 4days revealed that the presence of fibrils in the uncross-linked aAβ-UV sample, while the cross-linked aAβ+UV showed oligomers and a reduced number of fibrils. A minimum of three independent experiments was repeated to ensure the reproducibility of the findings.

**Figure 7.**
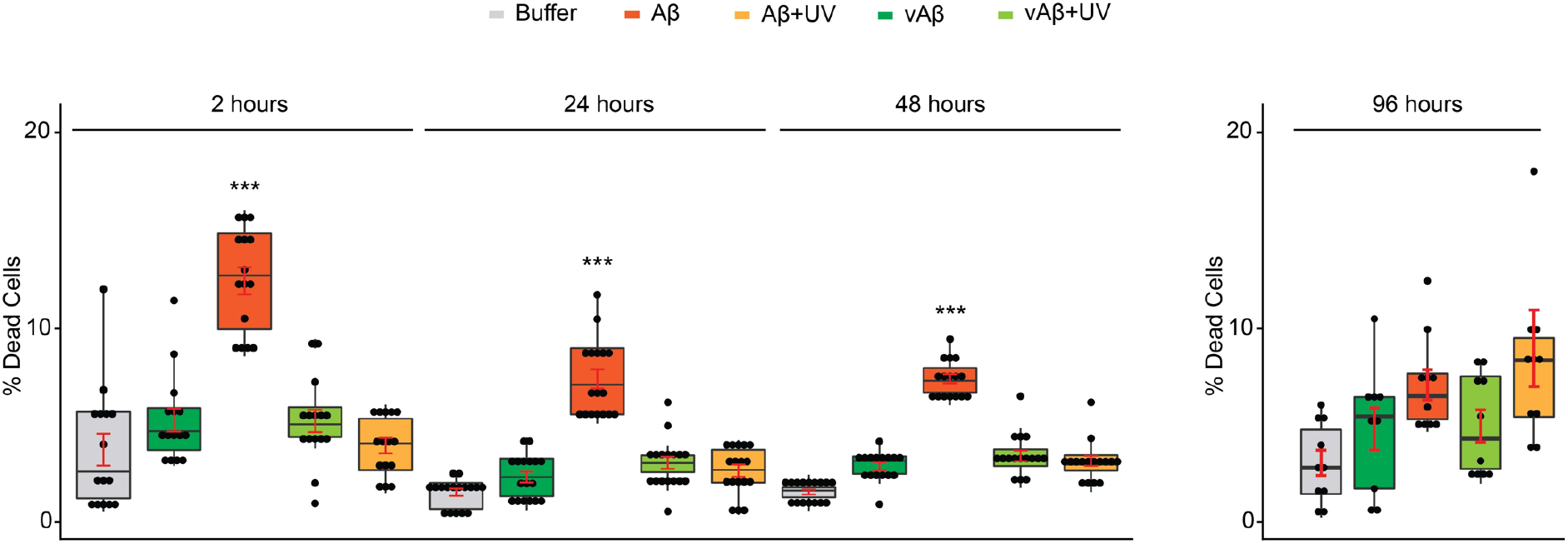
DiY-stabilised Aβ assemblies are not toxic to cells. Differentiated SH-SY5Y neuroblastoma cells were incubated with UV cross-linked or uncross-linked vAβ or Aβ for 3days, following which the percentage of dead cells was quantified using ReadyProbes Live/Dead Assay. The Aβ and vAβ samples were freshly prepared and DiY-stabilised with UV for 2h (Aβ+UV/vAβ+UV), or DiY-stabilised for 2h followed by a further incubation on bench and in the dark for 24h, 48h or 96h, before being administered to cells. vAβ and Aβ samples not exposed to UV were used as reference. Only the uncross-linked/assembling wild-type Aβ induced significant cell death. Experiments were repeated five times. *** P < 0.001.

### Self-assembly is important for Aβ toxicity

Multiple pieces of evidence have shown the detrimental role of Aβ on neuronal function that eventually leads to neuronal death^2–6^. Our Th-T, dot blot and TEM data showed that Aβ42 monomers self-assemble to form significant level of oligomers at two hours, and eventually protofibrils, fibrils and a network of fibrils at 4 days (Fig. 2, 6). Hence, the rate of self-assembly is high at early timepoints and plateaus at later timepoints when Aβ forms fibril network and plaques^3^. To investigate the role of self-assembly in toxicity, ReadyProbes live/dead assay was conducted on terminally differentiated SH-SY5Y neuroblastoma cells incubated for three days with wild-type Aβ following incubation for 2h, 24h, 48h and 96h to form oligomers, protofibril mixture and fibrils and then treated for two hours with UV (Aβ+UV) to induce DiY, or untreated (Aβ-UV). vAb was used as an assembly incompetent control^3^. Our results reveal a significantly higher level of cell death in cells incubated with Aβ after 2h of preparation, compared to cells treated with Aβ after 24h, 48h and 96h incubation as expected^3^. However, the Aβ samples stabilised with DiY showed no toxic effect to the cells, similar to the assembly incompetent cross-linked and non-cross-linked vAβ. This suggests a key role for self-assembly in the toxicity of Aβ.

## Discussion

In AD, Aβ self-assembles to dimers, oligomers, fibrils and eventually amyloid plaques – one of the key hallmarks of the disease. Aβ is generally accepted to play a key role in AD, but the mechanism behind its toxicity is still not completely understood. Numerous studies have searched for the elusive “toxic” species of and tried to characterise its structure and this has revealed supposedly toxic assemblies such as dimers, 12mers, *56 KDa, hexamers^28^. Here we show that DiY cross-linking significantly slows, or halfs the self-assembly of Aβ42 trapping it in a specific state. Here we compared the self-assembly and toxicity of assembly incompetent vAβ with wild type Aβ and DiY cross-linked intermediates.

Previous studies have suggested that DiY cross-linking can facilitate Aβ^29–32^, or inhibit or slow Aβ self-assembly^23,24^. Importantly, it has been shown that copper influences self-assembly differentially depending on the concentration and ratio^33,34^ which may go some way to explaining these discrepancies. Here we have explored this question by comparing oxidation of wild-type Aβ with vAβ which does not self-assemble^3^. MCO and photo-induced oxidation of vAβ revealed that DiY forms rapidly as early as 5 min post-oxidation, however, cross-linking did not lead to vAβ assembly even after 5 days in the oxidative environment or post-oxidation. This demonstrates that the formation of DiY doesn’t induce aggregation of the vAβ. DiY was also rapidly induced in wild-type Aβ using both MCO and photo-induced oxidation, however further assembly is inhibited. Co-incubation of cross-linked Aβ and freshly prepared uncrosslinked Aβ demonstrated significantly reduced assembly. This suggests that DiY cross-linking does not induce or facilitate the aggregation of the wild-type Aβ. Instead, it stabilises or traps Aβ assemblies and delay further elongation. This is supported by previous reports that showed that DiY cross-linked Aβ are slow to fibrilise and form long-lived soluble oligomeric aggregates^19–22^. Mass-spectrometry studies have revealed that DiY cross-linking leads to the stabilisation of Aβ40 in compact oligomeric species^22^, which is in strong support of our findings.

Aβ exists in a pool of monomers, soluble oligomers, and insoluble fibrils. Multiple studies have reported that the soluble Aβ oligomers in AD are comprised of dimers, trimers, tetramers, pentamers and decamers, Aβ-derived diffusible ligands (ADDLs), dodecamers, and Aβ*56^28^. A question therefore arises as to how DiY cross-linking impacts on these pool of assemblies. Our previous data showed that within 2h of preparation, Aβ exists mostly as oligomers with a random coil conformation^3^. Here, our results showed that the very slow oxidation induced by CuCl_2_ alone first facilitates the formation of Aβ oligomers followed by DiY cross-linking of the oligomers into stabilised oligomeric population. The more rapid MCO reaction results in DiY formation as early as 5 min post-oxidation whereby the Aβ becomes trapped in a pre-oligomeric conformation (as assessed using the antibody NU1). Th-T fluorescence, CD and TEM showed UV-induced cross-linking of early Aβ assemblies traps Aβ in a random coil confirmation and prevents or significantly delays further assembly into amyloid fibrils. UV-induced DiY cross-linking in preformed oligomer/protofibril assemblies, similarly results in the stabilisation this state and significantly delays further elongation to fibrils. Taken together, these results suggest that the timing of DiY formation in Aβ critically influences its assembly, leading to the stabilisation or significantly reduced assembly of the Aβ assemblies at the time of cross-linking.

Aβ self-assembly is believed to be important for its toxicity^3^ and many studies have implicated the role of oligomeric species in cytotoxic effects^35–37^. Here, we compared toxicity of the uncross-linked Aβ with DiY cross-linked Aβ whereby specific species in the assembly pathway have been stabilised. Aβ42 was DiY crossed link at different time points to stabilise a series of pre-oligomer, oligomeric, protofibrillar and fibrillar forms. We show that all these species are unable to induce cell death following three days of incubation with differentiated neuroblastoma cells. This finding is in conflict with previous studies that showed that DiY Aβ42 assemblies are toxic to cells^17^; DiY cross-linked Aβ40 dimers induce cell viability loss^19^ and that Th-T positive, DiY cross-linked Aβ40 fibrils were able to inhibit long-term potentiation (LTP)^20^. The observation that DiY vAβ is not toxic to cells supports our conclusions, suggesting that the presence of DiY alone is not sufficient to induce toxicity. The DiY crosslinked Aβ42 assemblies produced here are different from DiY Aβ40 reported by others^19,20^, as the DiY Aβ42 produced here loses Th-T binding, and does not proceed to form fibrils. However, we do not rule out the possibility that DiY Aβ preparation in our study and others also results in other oxidative modifications. Here we show that oxidation that leads to formation of DiY halts Aβ self-assembly and prevents cytotoxic effects. We therefore conclude that continued self-assembly is important for Aβ toxicity. We have previously demonstrated that assembly incompetent vAβ is not toxic to cells [3]. Our work also supports a role for DiY formation in the stability of Aβ42 species, which may be important for the formation of insoluble Aβ42 assemblies. We believe that the timing of the DiY formation may be critical. Formation of DiY on Aβ fibrils would promote its stability and formation of amyloid plaques. Indeed, we have previously shown the presence of DiY on Aβ plaques in AD brain tissue and demonstrated that DiY Aβ fibrils become highly insoluble and resistant to formic acid denaturation^15^.

In conclusion, DiY cross-linking specifically appears to promote the stabilisation, and not induce or facilitate, Aβ assembly. Our findings strongly suggest a role for self-assembly for Aβ toxicity. We show that Aβ exerts a high level of toxicity at a stage when self-assembly is high, compared to when the self-assembly rate is significantly diminished or abolished. This is observed even for those preparations where oligomeric Aβ has been stablised. Our work implies that the timing of DiY formation plays a key role in further assembly and stability of Aβ.

## Acknowledgements

This work was supported by funding from Alzheimer’s Society [345 (AS-PG-16b-010)] awarded to LCS and funding MBM.

## Author contributions

MBM planned and carried out the work. GB, YA and KM contributed experimental work. MBM and LCS wrote the paper. YA edited the paper. LCS managed the project.

